# Lifespan Oscillatory Dynamics in Lexical Production: A Population-based MEG Resting-State Analysis

**DOI:** 10.1101/2024.07.28.605484

**Authors:** Clément Guichet, Sylvain Harquel, Sophie Achard, Martial Mermillod, Monica Baciu

## Abstract

Lexical production remains relatively preserved across the lifespan, but cognitive control demands increase with age to support efficient semantic access. It suggests a domain-general and a language-specific component. Current neurocognitive models suggest the Default Mode Network (DMN) may drive the interplay between these components, impacting the trajectory of production performance with a pivotal shift around midlife. However, the corresponding time-varying architecture still needs clarification. Here, we leveraged MEG resting-state data from healthy adults aged 18-88 from a CamCAN population-based sample. We found that DMN temporal dynamics shift from anterior-ventral to posterior-dorsal states until midlife to mitigate word-finding challenges. Similarly, sensorimotor integration along this posterior path enhances cross-talk with lower-level circuitry as the dynamic information flow with more anterior, higher-order cognitive states gets compromised. It suggests a bottom-up, exploitation-based form of cognitive control in the aging brain, highlighting the interplay between abstraction, control, and perceptive-motor systems in preserving lexical production.

**Highlights:**

- Midlife is a pivotal period for time-varying functional connectivity
- DMN activation and deactivation drive the resting-state oscillatory architecture
- Enhanced posterior DMN temporal dynamics mitigates lexical production decline

## 1. Introduction

The increase in life expectancy is associated with a stringent need to understand the contributing factors of healthy and “successful” aging ^1^. Research in cognitive aging has uncovered significant insights into the natural progression of cognitive functions with age. While several cognitive abilities, including episodic and working memory, attentional and inhibitory control, begin to decline in early adulthood ^2^, language skills remain relatively preserved ^3,4^. Given the fundamental role of language in human cognition ^5,6^, gaining a deeper understanding of the neural mechanisms that support the maintenance of language functions is essential.

Relatedly, studies suggest that language interacts across various cognitive domains, potentially as the common thread behind neurocognitive changes supported by an intricate interplay of perceptive-motor and cognitive systems ^7–10^. For example, the Language-union-Memory (LLM) framework ^11^ proposes that language processing involves the bidirectional flow of information between external sensory modalities and long-term memory, flexibly monitored by task-relevant control processes. Evidence also shows that the language connectome ^12,13^ engages core regions ^14^ alongside large-scale brain networks reflecting control-executive, abstract-conceptual ^15^, and sensorimotor processing.

Changes in hemodynamic activity within these networks have allowed for a comprehensive description of the mechanisms involved in preserving language function across the lifespan. For example, the Lexical Access and Retrieval in Aging or LARA model ^16,17^ posits domain-general and language-specific compensatory mechanisms, reflecting the interplay between executive functioning and semantic memory. In this context, longer naming latencies and more frequent tip-of-the-tongue situations with age could be mitigated by enhanced semantic access thanks to a distributed anatomo-functional reorganization ^12,18–20^. Specifically, the enrichment of semantic repositories ^2,21^ could represent a strategy for maintaining the retrieval of words with stronger semantic connections ^22^ and conceptual categorization ^23,24^ contingent upon accumulated vocabulary knowledge throughout adulthood ^25,26^. However, more pronounced inhibition deficits could challenge the ability to ignore task-irrelevant information ^27–29^, leading to longer searches through a growing semantic store. Indeed, reduced access and retrieval of lexico-semantic representations ^3,30^ may induce more automatic and distracting thoughts, leading to a global decline in selecting the appropriate semantic information ^31–34^, with difficulties beginning around midlife ^4,22,35^.

At the brain level, the central role of semantic control in lexical production is reflected by activations within the multiple demand (MD) and default mode network (DMN). The MD network ^36^ is composed of a core set of regions from the fronto-parietal (FPN), dorso-attentional (DAN), and cingulo-opercular (CON) networks that are relevant for goal-directed cognition ^37–40^. The DMN, usually activated in the absence of external stimuli, plays a crucial role in self-directed and semantically rich cognitive processes ^41–43^. Importantly, semantic control regions appear to adjust their activation to cognitively demanding semantic processes by engaging a set of areas “sandwiched” between the MD and DMN ^44^. This aligns with current models ^45,46^ delineating a control subsystem involving the anterior-ventral parts of the IFG ^47^ and pMTG ^33,48^ and a language-specific subsystem primarily storing multimodal representations in the ventral ATL ^49,50^.

In the context of healthy aging, models such as the Default-Executive-Coupling Hypothesis of Aging (DECHA) ^29^ propose that lateral PFC engagement within the FPN is functionally coupled with DMN suppression, thus integrating domain-general and language-specific mechanisms central to semantic control. Evidence supports that DMN suppression optimizes goal-driven executive tasks, including semantic classification ^51^, by reducing self-directed processes such as mind-wandering ^52–55^. In older ages, reduced DMN suppression may disrupt the coupling with FPN regions, leading to poorer performances when prior semantic knowledge is irrelevant to the current task. For example, increased anterior DMN connectivity (right ATL-mPFC) in older adults has been found to strengthen verbal semantics when prior knowledge is congruent with the task but impairs the ability to discern complex semantic relationships that require more top-down control ^56^. Recently, a study introduced a new framework that redefines the DMN-FPN coupling through the lens of brain homeostasis (see the SENECA model^12^). The model suggests that DMN suppression offsets the cost of recruiting additional control resources until midlife, efficiently maintaining highly integrative and “energy-costly” functions such as lexical production. In older ages, however, reduced DMN suppression may no longer offset this cost, prompting a switch towards a more “energy-efficient” strategy based on exploiting memory-guided control inputs, with enhanced cross-talk with the CON network ^57,58^ rather than the FPN.

While time-averaged properties of these mechanisms have been investigated using fMRI, studies examining moment-to-moment inter-regional synchrony, or dynamic FC, remain more scarce. In that regard, magnetoencephalography (MEG) offers a higher temporal resolution with comparable resting-state functional connectivity, as measured with fMRI ^59^. Therefore, the main goal of this study was to provide an integrated spatial, temporal, and spectral description of the mechanisms involved in maintaining language function across the lifespan, specifically considering the time-varying dynamics of canonical resting-state networks.

Overall, we hypothesized that DMN oscillatory activity will drive the dynamic resting-state architecture, maintaining lexical production performance according to two mechanisms: (i) a Domain-General (DG) mechanism engaging bilateral prefrontal cortices correlated with flexible abilities (i.e., problem-solving). We primarily expected changes in alpha band synchrony, reflecting greater top-down control demands for semantic word retrieval ^60^ in older adults. We also expected shorter and more infrequent DMN-FPN coupling beyond midlife, reflecting challenges in dynamically modulating their connectivity ^29^. (ii) a Language-Specific (LS) mechanism supporting access to lexico-semantic representations within lateral and mesial left-lateralized frontotemporal networks ^17^. We expected this mechanism to peak in midlife, reflecting heightened information flow in brain states implementing DMN-CON coupling ^12^.

## 2. Results

We leveraged MEG resting-state data from healthy adults aged 18-88 from the CamCAN population-based cohort ^61^ and assessed lexical production performance using 8 direct or indirect cognitive measures. The study analysis pipeline is completely unsupervised and data-driven: (i) First, we identified dynamic brain states common to all individuals using the Hidden Markov Model (HMM); (ii) Second, we related these states’ spatial, temporal, and spectral features to lexical production performance across the lifespan. Figure 1 presents an overview of the pipeline.

**Figure 1.**
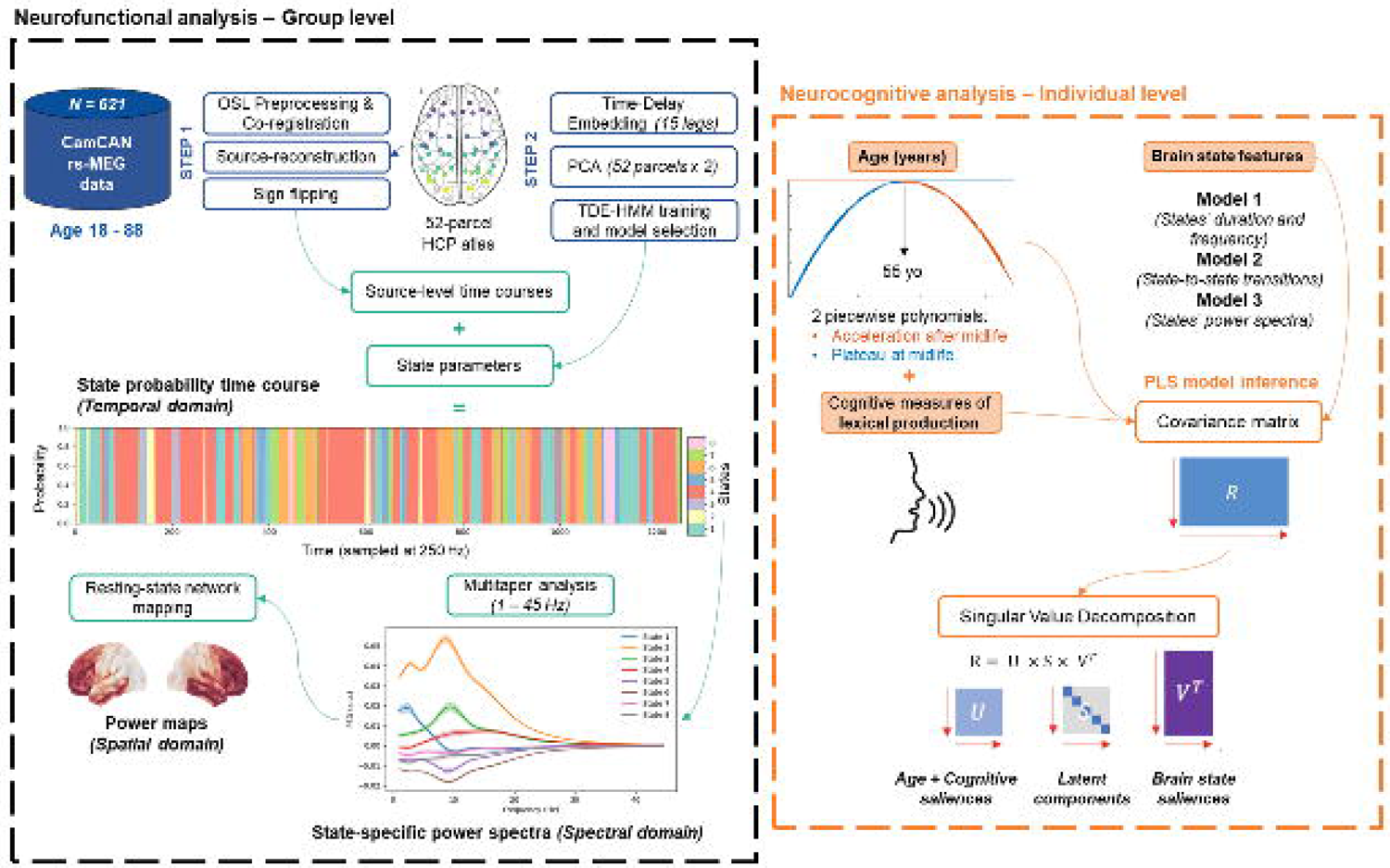
Overview of the analysis pipeline. *(Neurofunctional).* Following the preprocessing steps, we employed the Time-Delay Embedded Hidden Markov Model (TDE-HMM) to identify functional brain states from resting-state MEG time series data. We then extracted each state’s group-level temporal, spectral, and spatial features *(Neurocognitive).* We examined the relationship between age-related lexical production performance and brain state features using the Partial Least Squares (PLS) technique. Age was parametrized using 2 piecewise polynomials centered at 55 years old to explore non-linear trajectories during inference. Cognitive performance was assessed with 8 tasks directly or indirectly related to lexical production. We further considered 3 types of brain features in 3 distinct models: temporal metrics (duration & frequency), transition probabilities from one state to another, and the power spectra of the top 20 state-relevant parcels. For details, please refer to the Method section.

### 2.1. Dynamic brain states

Leveraging the Hidden Markov Model approach, we modeled resting-state MEG activity and identified eight brain states whose activations wax and wane over time. Consistent with our hypothesis, we observed that DMN activity and suppression are crucial elements of time-varying functional activity. As shown in Figure 2, we observed that they are spectrally coupled within the 1-8 Hz and 1-25 Hz frequency range, respectively, in the anterior-ventral axis (State 1-7), where the DMN preferentially binds with the FPN network, and posterior-dorsal axis (State 2-8) where the DMN binds with lower-level circuitry. Below, we describe the group-level spatio-spectral features of each state, beginning with the states that show the largest oscillatory changes compared to the time average (see Figure S1-2, Appendix S2).

**Figure 2.**
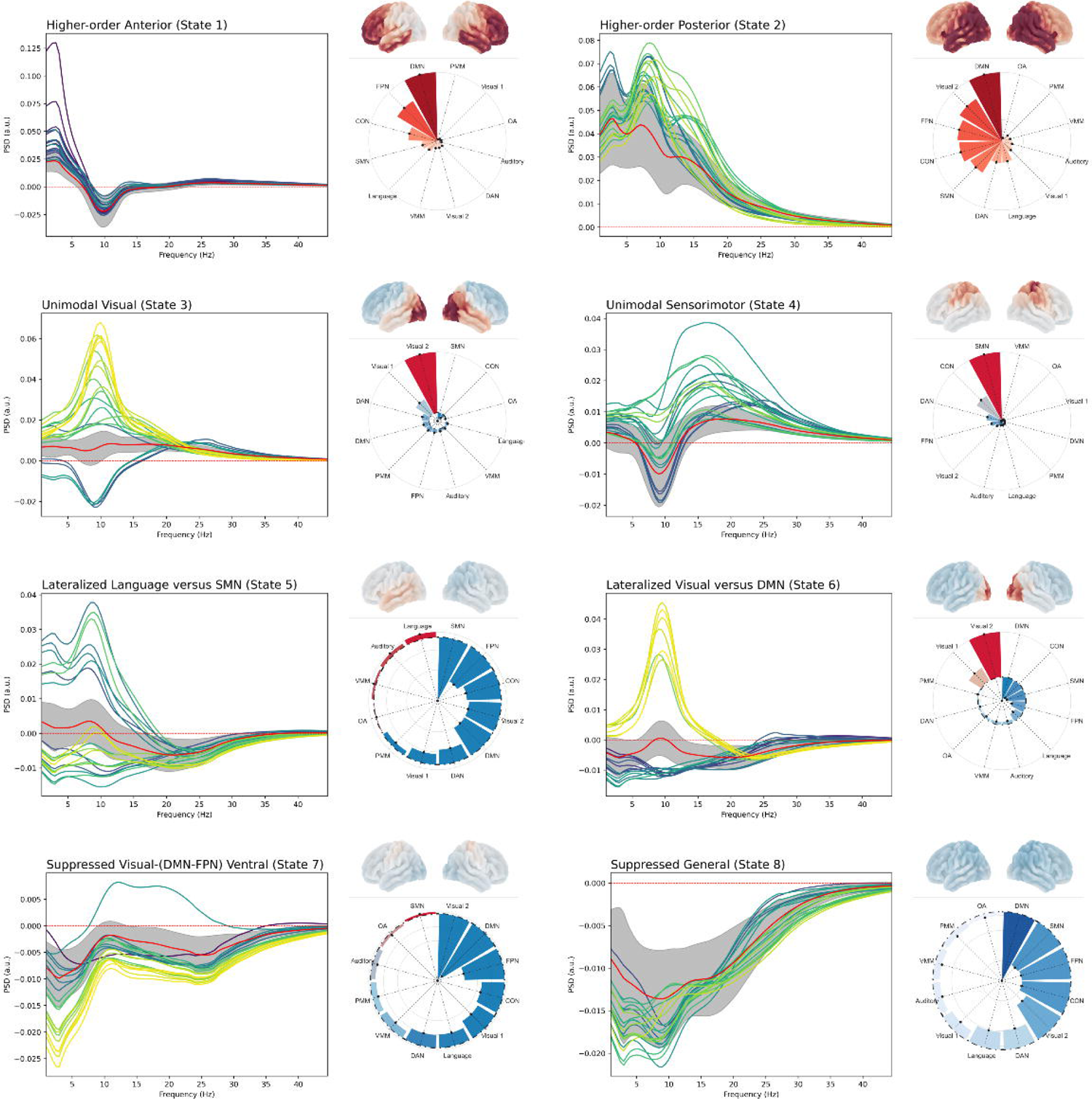
Dynamic spatio-spectral content of the 8 brain states. Main plot. Illustrates the power spectra of the 20 channels with the largest departure from static activity, thus reflecting the dynamic features of each state. The colors of each channel follow their spatial location (yellow = posterior, green = middle, purple = anterior). The red line represents the mean power across all channels and participants. The grey strip is the variance among participants**. Brain plot.** The group-level spectrum of each state is integrated into the spatial domain (red = activation; blue = suppression). **Radar plot.** Resting-state network topography (red = positive contribution; blue = negative contribution). For additional details, see Table S1-2-3, Appendix S3. *Network abbreviations: DMN (Default Mode), FPN (Fronto-Parietal), CON (Cingulo-opercular), DAN (Dorso-Attentional), SMN (Sensorimotor), VMM/PMM (Ventro-/Posterior Multimodal), OA (Orbito-Olfactive)*.

#### Higher-order states

State 1 implies high activity in the delta/theta band (∼1-7 Hz) combined with a small release of alpha power (∼8-12 Hz). This state engages medial and lateral frontal areas of the DMN and FPN networks, respectively. In comparison, State 2 involves an increase in oscillatory activity across a wider 1-25 Hz frequency range, with prominent bursts of activity in the delta and alpha bands. This state engages temporoparietal areas primarily in the lateral and posterior parts of the DMN, FPN, and CON networks. Interestingly, State 2 shows greater variability among individuals, which could reflect the greater effect of age on this state.

#### Unimodal states

State 3 is an alpha-band powered state with increased oscillatory activity in visuo-occipital areas. In comparison, State 4 shows increased beta-band pre-/post-central sensorimotor activity combined with a release of alpha power in more anterior regions. This activity release also shows ample individual variability, suggesting an age-dependent modulation.

#### Lateralized states

States 5 and 6 are characterized by opposite oscillatory activity between the two hemispheres, showing mild positive or negative changes in the alpha band. State 5 engages left-hemispheric language-auditory circuitry at the expense of dense right-hemispheric sensorimotor-visual activity. In contrast, state 6 engages right-dominant visuo-occipital circuitry at the expense of left temporal DMN activity.

#### Suppressed states

States 7 and 8 exhibit a mild suppression of activity relative to the average over time. State 7 specifically engages more ventral posterior regions with the same network and spectral characteristics as State 1. Similarly, State 8 is more spatially distributed across a wider 1-25 Hz frequency range, mirroring the network and spectral characteristics of State 2.

### 2.2. Neurocognitive analyses

Using the Partial Least Squares (PLS) analysis, we sought to uncover how brain state features relate to changes in lexical production performance across the lifespan. We specifically examine temporal characteristics (duration and frequency of state activations), transitions between states, and spectral characteristics.

In line with our hypothesis, we consistently found (i) a **domain-general component** denoting an acceleration in cognitive control deficits beyond midlife and primarily underpinned by changes in temporal dynamics (see Figure 3A-B).; (ii) a **semantic-specific component** denoting an inverted U-shaped trajectory across the lifespan, associated with enhanced semantic access and spectral changes (see Figure 3C-D). Model diagnostics are reported in Figure S6, Appendix S2.

**Figure 3.**
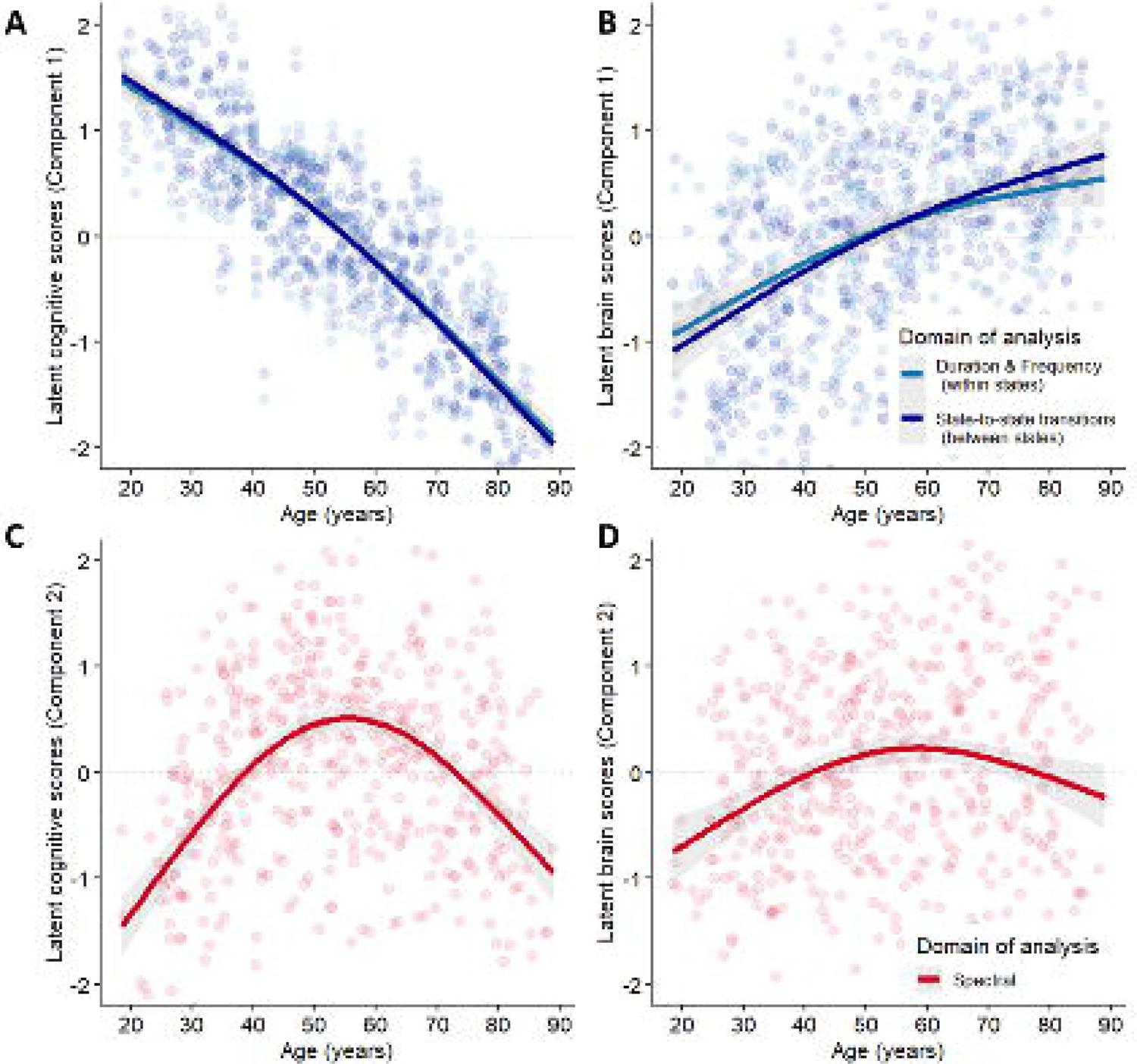
Latent age-related trajectories. Projection of the latent age-related scores associated with the cognitive and brain latent variables. **A-B.** The first component correlates with the cognitive control aspects of lexical production and shows accelerated cognitive decline beyond age 60 in the temporal models. **C-D.** The second component correlates with the semantic/multitasking processes related to lexical production and peaks in midlife. Only the spectral model explains a significant amount of variance. The inflection point was fixed a priori to 55 years old (see Method).

#### 2.2.1. Component 1 – Temporal features reflect domain-general abilities

Figure 3A shows that the first component negatively correlates with age, denoting more pronounced difficulties beyond age 60. This may primarily affect performances in tasks assessing fluid intelligence and picture-naming abilities and could be exacerbated by low education attainment (Table 2).

**Table 1.** Neuropsychological tasks and associated cognitive processes were considered in this study. A detailed description can be found in Appendix S1. LP=Language production.

| Neuropsychological test | Cognitive processes | Task type and relation to lexical production |
| --- | --- | --- |
| Picture naming | Language (phonological and semantic access) | LP, Direct |
| Verbal fluency | Language (phonological and semantic access) and EF | LP, Direct |
| Tip-of-the-tongue situations | Language (retrieval) and EF (monitoring) | LP, Direct |
| Proverb comprehension | Language (comprehension) and EF (abstraction) | Language, Indirect |
| Sentence comprehension | Language (syntax, semantic) | Language, Indirect |
| Story recall | Long-term memory | Language, Indirect |
| Cattell task | EF (Logical reasoning, problem-solving) | Domain-general, Indirect |
| Hotel task | EF (multitasking, planification, self-monitoring) | Domain-general, Indirect |

**Table 2.**
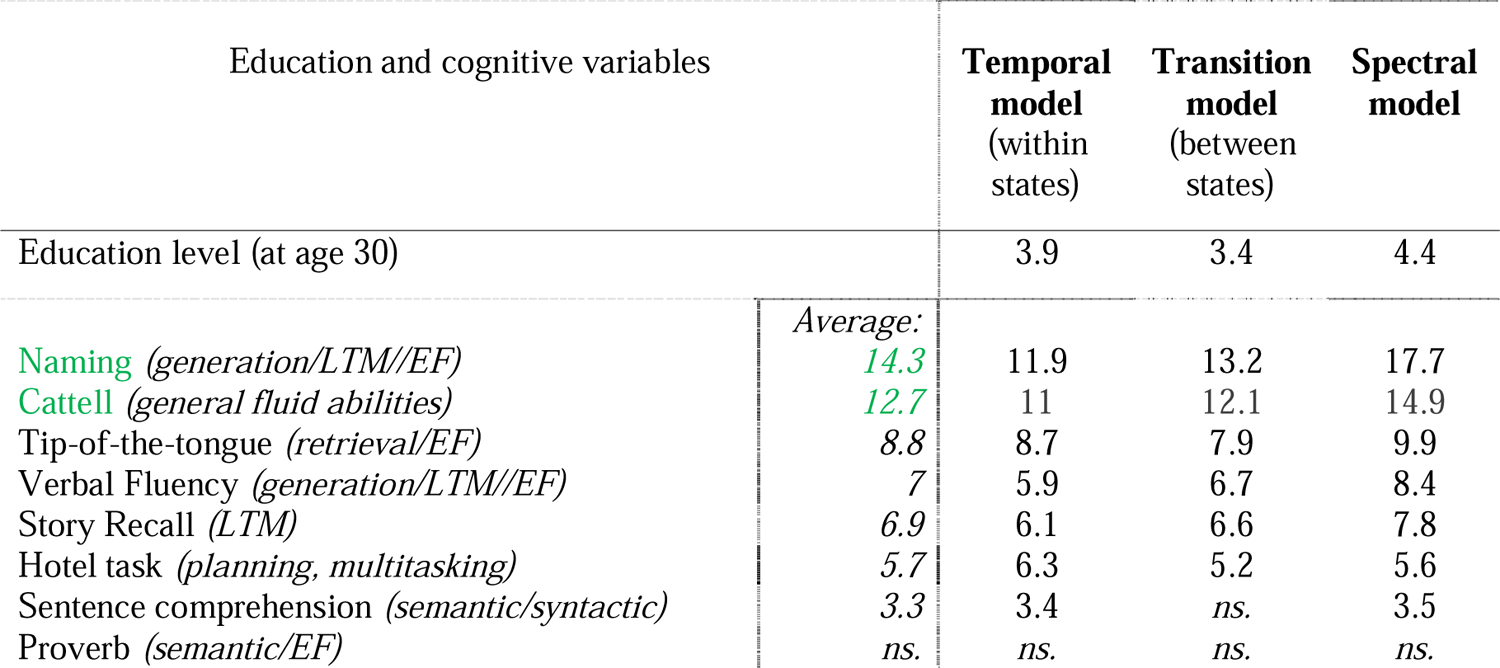
Bootstrap sampling ratios (BSR) of education and cognitive variables for LC1. Only salient (±3) bootstrap sampling ratios (BSR) are presented. *Abbreviation: ns. (not significant)*.

Interestingly, temporal changes within and between brain states explained most of the total shared variance (86.93% and 84.3%; *p*_FDR_ < .001) compared to the spectral model (58.75%; *p*_FDR_ < .001). This suggests that temporal dynamics adequately capture the lifespan trajectory of cognitive control performances involved in lexical production. Moreover, we note that some temporal changes may work to mitigate control deficits as a reduction in these changes beyond midlife concurs with an acceleration in cognitive decline (see Figure 3A). Thus, we primarily focus on the salient temporal features associated with this first component. Results on the spectral model are available in Appendix S2.

#### Duration and frequency (within states)

Figure 4 shows two main patterns regarding the duration and frequency of state activations contributing to the trajectories shown in Figure 3B: the first pattern involves states engaging DMN regions, and the second pattern involves SMN regions.

**Figure 4.**
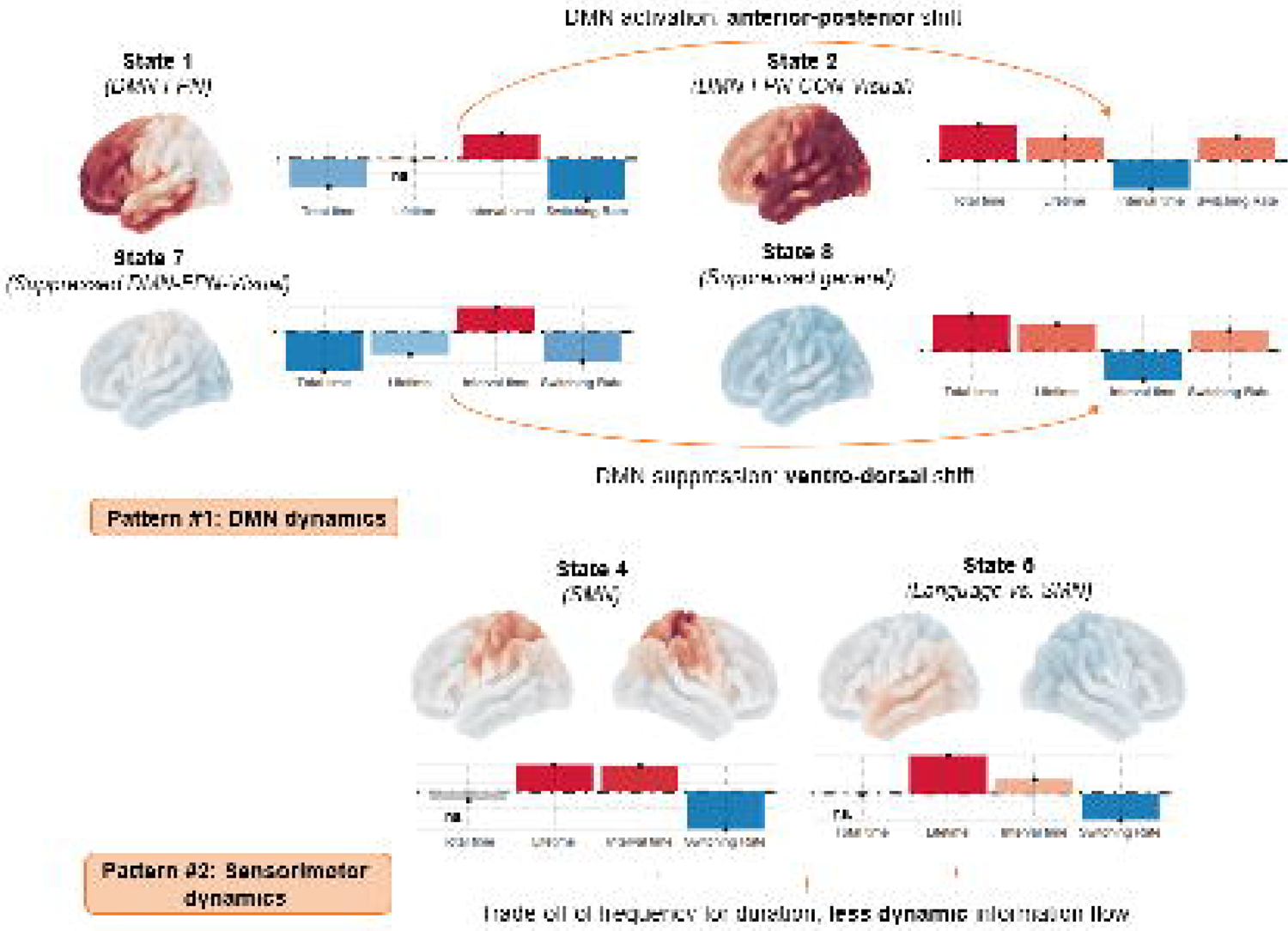
Bootstrap sampling ratios (BSR) for temporal metrics. We report two patterns contributing to the lifespan trajectory shown in Figure 3B. The first pattern represents an anterior-posterior DMN activity and ventral-dorsal DMN suppression shift. The second pattern represents a reduction of dynamic activity in sensorimotor cortices. **Brain plot.** The group-level spectrum of each state is integrated into the spatial domain (red = activation; blue = suppression). *ns. (not significant); Network abbreviations: DMN (Default Mode), FPN (Fronto-Parietal), CON (Cingulo-opercular), SMN (Sensorimotor)*.

More specifically, the first pattern is characterized by an increase in overall time spent activating the states (state 2; FO = 11.5; state 8; FO = 11.7) where the DMN binds with both the attentional (FPN, CON) and sensory-visual networks (SMN, Visual2). This increase reflects shorter interval times between activations (INT = −9.3/-9.3), more frequent state visits (SR = 7.6/6.5), and longer activations (LT = 7.6/8.8). Interestingly, this covaries with decreased overall time spent activating the states (state 1; FO = −12.1; state 7; FO = −16.6) where the DMN binds with the FPN and Visual network, respectively, in prefrontal and ventral posterior areas. This decrease reflects longer interval times (INT = 10.7/10.4) and less frequent state visits (SR = −17.7/-12.6), but no significant changes in mean lifetime when the state is active (see also Table S1, Appendix S2 for a summary).

The second pattern explicitly involves the sensorimotor network. Indeed, we observed that state 4 and state 5 have more sustained activations (LT = 6.9/12.7) but less frequent visits (SR = −9.6/-8.8) and longer interval times between state visits (INT = 6.6/4.7). Given that the overall time spent in these states does not change significantly, this alteration may be interpreted as a trade-off between the frequency and duration of activation.

In summary, our findings suggest an anterior-to-posterior and ventral-to-dorsal shift in the brain states’ temporal dynamics, all engaging the DMN. State activations on the anterior (state 1) and ventral (state 7) ends get more infrequent with age (although not shorter), whereas state activations on the posterior (state 2) and dorsal (state 8) ends get both more frequent and longer. Moreover, it seems that sensorimotor activation may become more inflexible with advancing age as the duration of state activations increases, but their frequency decreases. A thorough interpretation of these patterns is proposed in the discussion section.

#### State-to-state transitions (between states)

When considering successive state-to-state transitions, our results revealed a similar anterior-posterior and ventral-dorsal shift in how information flows is re-orchestrated, explicitly highlighting critical alterations in sensorimotor processing.

On the anterior and ventral ends, Figure 5 shows reduced transitions (i) anteriorly towards the higher-order state, especially after the sensorimotor regions were active (transition state 4 to 1 = −19), and (ii) ventrally after a release in sensorimotor oscillatory activity (transition state 5 to 7 = −14). More generally, this reflects reduced information flow from higher-order states to the sensorimotor state (states 1 & 2 to state 4 = −7.9/-7.5). This is paralleled by preferential transitions from the sensorimotor state towards heightened alpha-band activity in visuo-occipital areas (state 4 to 3 = 6.8, state 4 to 6 = 11), thus suggesting that sensorimotor flows more posteriorly with age. Relatedly, we observed more frequent transitions towards the states on the posterior and dorsal ends, especially after a release in sensorimotor oscillatory activity (state 5 to 2 = 8.1; state 5 to 8 = 9.8) or after a modulation of alpha-band visuo-occipital activity (state 3 to 2 = 11; state 6 to state 8 = 14). Similarly, we note reduced information flow along the dorso-ventral axis between the suppressed states (state 8 to 7 = −13), in line with the previous observation that DMN suppression preferentially diffuses within dorso-posterior areas with age.

**Figure 5.**
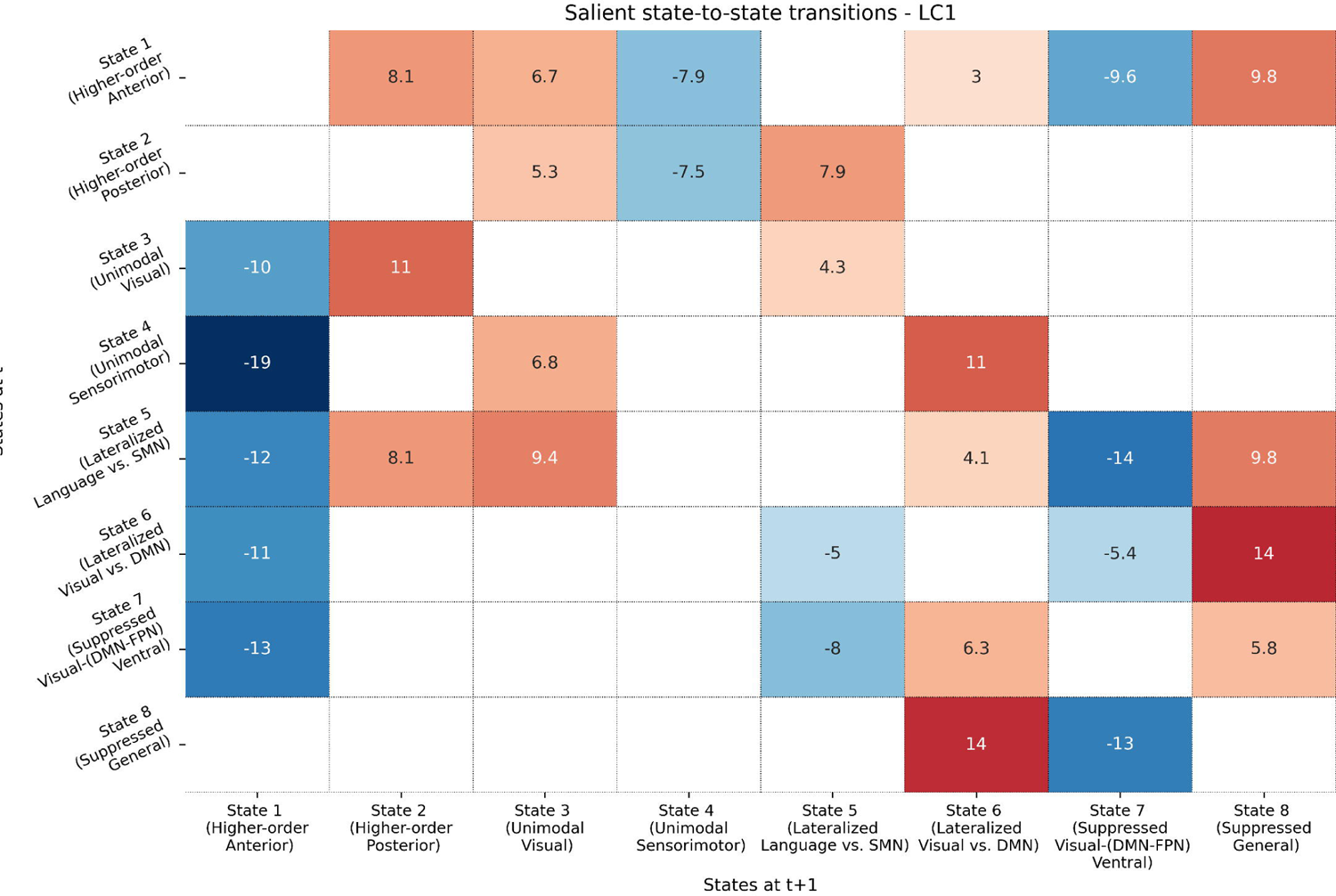
Bootstrap sampling ratios (BSR) for state-to-state transitions. Each cell corresponds to state-to-state transitions from time t to t+1. For example, the transition from the unimodal sensorimotor state (on the y-axis) to the higher-order anterior state (on the x-axis) is significantly reduced as age increases (BSR = −19). In other words, this transition is negatively correlated to the trajectory highlighted in Figure 3B.

#### 2.2.2. Component 2 – Spectral features reflect semantic-specific processes

The second latent component was significant across all models, but only spectral information explained a significant portion of the variance (28.68%; *p*_FDR_ < .001). This component captured semantic aspects of lexical production such as semantic abstraction (BSR_proverb task_ = 10) and comprehension (BSR = 4) but also highlighted the multitasking/self-monitoring processes, as measured by the Hotel Task (BSR_hotel task_ = 9.2), that could be recruited for maintaining verbal fluency (BSR = 9.2) and naming abilities (BSR = 5.7). We also note better long-term memory performances (BSR = 4.3), which continue to suggest the exploitation of stored knowledge in semantic memory for lexical production. Figure 6 shows that this cognitive outcome peaks in midlife and positively correlates with education attainment (BSR = 5.2). At the brain level, this mapped onto a general increase in the upper part of the spectrum (>13 Hz) across all states and a release of activity in the lower part (<13 Hz).

**Figure 6.**
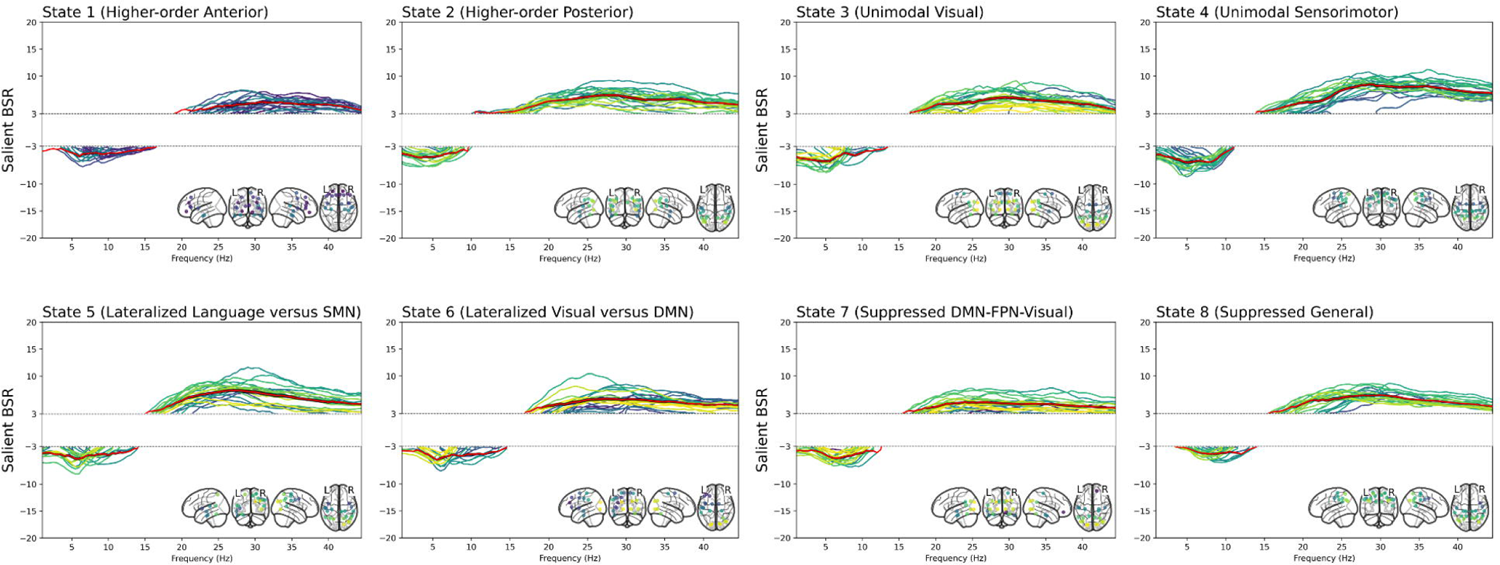
Bootstrap sampling ratios (BSR) in the spectral domain. The colors of each channel follow their spatial location (yellow = posterior, green = middle, purple = anterior). Red is the mean salient BSR value; the grey strip is the standard error. The white band in the range y = −3 to y = 3 masks BSR values below the significance threshold.

## 3. Discussion

As individuals age, lexical production, including word generation and retrieval, may increasingly demand cognitive control to manipulate and access the semantic store efficiently^49^. While semantic repositories significantly expand with age, cognitive control decline may lead to longer naming latencies, typically beginning around midlife. Thus, the present study aims to understand the neurocognitive mechanisms that delay the onset of lexical production decline with age, focusing on the role of semantic control in preserving language function.

Research has primarily utilized imaging techniques with low temporal resolution, such as fMRI, to investigate these mechanisms ^18,34^. Consequently, more is needed to understand the brain-wide changes in spontaneous oscillatory dynamics accompanying the onset of lexical production decline across the lifespan. To address this gap, our study leveraged recent machine learning advances for modeling the magnetoencephalographic (MEG) resting-state activity of 621 participants aged 18-88 from the CamCAN cohort ^61^.

Importantly, our study reaffirms that midlife is a turning point for spontaneous network oscillatory dynamics underlying language performance, consistent with previous MEG ^62,63^, time-averaged fMRI ^12^, and diffusion-weighted imaging (DWI)^19,64^. This further underscores the importance of studying the middle-aged brain from a multimodal perspective ^65,66^ as this period of life could be “prognostic of future cognitive outcomes” ^67^.

### DMN activation and suppression are spatially and spectrally coupled

From a neurofunctional perspective, we identified recurrent dynamic networks or *states* at the group level, providing an integrated spatial, temporal, and spectral description of the brain’s functional dynamics at rest (Figure 2). While the spatial topographies of certain brain states are strongly similar to low-level fMRI resting-state networks, areas of the Default Mode Network (DMN) are distributed within four states, reflecting its synchronization with multiple networks across frequency bands.

Relatedly, our results align with previous studies reporting a dissociation along the anterior-posterior axis when DMN drives higher-order states’ activations ^68^ and the ventro-dorsal axis when DMN activity decreases alongside visuo-occipital processing ^69^. Specifically, in the delta/theta frequency range, anterior DMN-FPN activity straddling the prefrontal-temporal cortex (State 1) couples with DMN-FPN-Visual suppression in ventral occipital areas (State 7). In the wider 1-25 Hz frequency range, more integrated temporoparietal DMN activity with attentional and visuo-sensorimotor subsystems (State 2) is paralleled by spatially diffuse suppression in dorso-posterior areas of the same networks. Our study shows that spontaneous DMN activation and deactivation are spectrally coupled across the cortex. This further supports that DMN oscillations represent a distributed pattern of increased power aiding in the coordination of information flow across the brain ^68,70,71^.

### Enhanced dorso-posterior DMN temporal dynamics: bottom-up compensation?

From a neurocognitive perspective, changes in the temporal dynamics of these coupling patterns explained as much as 85% of the age-related trajectory associated with lexical production performance (Section 2.2), particularly considering the decline in fluid abilities, word generation, and retrieval. That is, anterior and ventral activity in States 1 and 7, where the DMN preferentially binds with the FPN, becomes less frequent with age, with fewer transitions to these states. In contrast, posterior and dorsal release of activity in States 2 and 8 show the opposite pattern. Consistent with our hypotheses, this suggests that DMN temporal dynamics are key to the age-related mechanisms underlying lexical production decline. This also demonstrates that temporal features best predict domain-general aspects of production, which aligns with the idea that inhibitory control is temporally focused ^72^. While the shift in spontaneous DMN activation/suppression orchestration strongly correlates with age-related cognitive decline, this could partially represent a compensatory mechanism.

Indeed, we found that cognitive decline accelerates when temporal changes plateau beyond midlife (refer to the latent trajectories in Figure 3A-B). Thus, compensatory and maladaptive mechanisms may coexist in the context of cognitive decline ^73^. In that regard, our correlational results alone cannot fully discern the temporal changes contributing to such compensation from those reflecting cognitive decline. For example, older adults’ sustained, spatially diffuse, and less frequency-specific dynamics in dorso-posterior states could compromise cognitive flexibility ^74^ and reflect the dedifferentiation process typically observed in fMRI ^75^. However, this could enhance the cross-talk between the DMN and lower-level circuitry in these states, representing a bottom-up strategy for preserving lexical production. In line with this, recent work suggests that increased dwell time in older adults could signal an exploitative search ^76^, perhaps via visually mediated semantic processes ^77^.

Previous studies may provide further evidence supporting a broader exploration-to-exploitation shift during healthy cognitive aging ^76,78^. According to this framework, older adults perform best when exploiting accumulated semantic knowledge, e.g., in inferential naming tasks that rely on semantic/episodic associations as opposed to traditional picture naming paradigms ^79^. This is especially relevant to our results, considering that increased recruitment of posterior regions has been reported in older adults for object naming ^80^ and semantic processing ^81^.

Moreover, the anterior-posterior shift in higher-order cognitive states (from State 1 to 2; Figure 4) echoes a similar spatio-functional dissociation in the form of control processes relevant for goal-directed behavior ^82^. Lateral and medial parts of the frontopolar cortex, as found in State 1, may facilitate goal selection and monitoring which authors associate with an exploration drive. In contrast, more posterior cortical regions may be geared towards implementing cognitive control to optimize the current task, which authors associate with an exploitation drive. Thus, the anterior-to-posterior shift in DMN temporal activity seems to have a strong cognitive justification, reflecting a shift from an exploration-driven to an exploitation-based strategy to support lexical production performance.

Although the metabolic underpinnings have not been addressed in this study, our results also corroborates with the predictions made by the SENECA model ^12^ in line with recent work showing that older adults have a smaller energy budget previously ^83^. Indeed, the additional engagement of the Cingulo-Opercular Network (CON) network in more posterior states (Section 2.2) supports the hypothesis that older adults gradually adopt a more “energy-efficient”, bottom-up form of cognitive control through increased transitions between the DMN (vmPFC & PCC) and the CON (fronto-insular and anterior cingulate cortices) ^57,78,84^.

### Sensorimotor integration is less dynamic and redirected along a posterior route with age

Another key observation is that sensorimotor integration becomes less dynamic with advancing age, trading off frequency for increased duration of state activations (Figure 4). Specifically, results from the state-to-state transition model suggest that older adults may struggle to dynamically modulate information flow with anterior, higher-order cognitive states that engage the DMN and FPN. This highlights that the cognitive control circuitry associated with word generation and retrieval likely depends on interactions with pre- and post-central areas of the sensorimotor network throughout the lifespan. These findings are consistent with claims that the sensorimotor network shapes the dynamic resting-state activity across the lifespan ^85,86^, which is critical in optimizing exploratory-driven goal selection ^87^.

Relatedly, sensorimotor information (States 4-5) was more often redirected along a posterior route via visuo-occipital areas (States 3-6) to dorso-posterior states where DMN converges with lower-level circuitry (States 2 and 8). Acute engagement of the visuo-occipital regions in the alpha-band may thus facilitate information flow between sensorimotor and dorso-posterior DMN in older adulthood, hypothetically lowering the metabolic demands of maintaining long-distance functional connections ^88^.

### Spontaneous low gamma-band activity gateways semantic memory-guided control inputs

More marginally, our model highlighted a semantic-specific component by faster oscillatory rhythmic activity (13 - 45Hz) in midlife and associated with enhanced semantic access and the multitasking aspect of control processes. This could highlight the enhanced semantic access, which relies on the bottom-up aspect of executive functioning suited to apply existing knowledge to real-life scenarios (i.e., exploitation-based mentation). Indeed, the Hotel task employed in this study appears to be a more ecologically valid assessment ^89^ than a standardized, distraction-free environment typically used to perform the Cattell task (i.e., to assess fluid reasoning). At the brain level, this is consistent with gamma power (30-150 Hz) being the main physiological basis of bottom-up information processing and could reflect the activation of local functional networks supporting semantic representations ^72,90^. However, this result warrants further investigation as low gamma-band activity in our study is limited to 45 Hz and may be subject to interpretation limitations inherent to time-delay embedding ^68,91^.

### Limitations of the study

Other methodological limitations include the use of cross-sectional data, which precludes direct inferences about the aging process, and the lack of cognitive reserve proxies beyond education levels. The Hidden Markov Model approach also holds certain assumptions, such as the a priori specification of the number of states and a Gaussian observation model, which may oversimplify the underlying network dynamics. Regarding the Partial Least Squares models, improving the fit between spectral information and cognitive outcomes may require examining cross-frequency couplings, given their crucial role in integrating, coordinating, and regulating neuronal activity ^92^. Additionally, we mention that we defined the turning point at age 55, fundamentally limiting the range of potential age trajectories explored during model inference. However, we argue that this approach arguably offers a less biased methodology for modeling age-related effects than discrete age groups. Accordingly, future work could fix the quadratic term at different ages following the study hypotheses or explore alternative unsupervised techniques better suited to explore nonlinear relationships with age. More complex models that relax Gaussianity and linearity assumptions, such as deep learning approaches, could provide additional insights. Future work should also explore how age-related changes reported in this study compare with task-related oscillatory dynamics.

## 4. Conclusion

This study aimed to elucidate the time-varying neurofunctional mechanisms involved in preserving lexical production across the lifespan. Our findings highlight two coupling patterns between DMN activation and suppression within distinct frequency bands: one along the anterior-ventral axis and another along the posterior-dorsal axis. To mitigate word-finding difficulties, the aging brain undergoes significant changes in how these patterns unfold at rest. When information flow with mediolateral prefrontal areas is compromised, older adults may preferentially engage posterior regions to enhance the cross-talk between the DMN and lower-level circuitry, indicating a greater reliance on a bottom-up, exploitation-based form of cognitive control. Our results suggest that this posterior route could facilitate information flow between sensorimotor processing and higher-level cognition through alpha-powered visuo-occipital activity. In sum, we propose that the onset of lexical production decline at midlife could reflect challenges in mobilizing this posterior circuitry to exploit semantic memory-guided control inputs. We recommend further examination of the interplay between abstraction, control, and action systems to fully capture the mechanisms involved in preserving lexical production.

## Acknowledgments

This work was supported by the ANR project ANR-15-IDEX-02 and MIAI @Grenoble Alpes (ANR-19-P3IA-0003). This project has received financial support from the CNRS through the MITI interdisciplinary programs. The Cambridge Centre for Ageing and Neuroscience (Cam-CAN) research was supported by the Biotechnology and Biological Sciences Research Council (grant number BB/H008217/1).

## Author contributions

Conceptualization: C.G. and M.B.; Methodology, Formal Analysis & Data Curation: C.G.; Writing – Original Draft: C.G. and M.B.; Writing – Review & Editing: C.G., S.H., S.A., M.M., and M.B.; Funding Acquisition: S.A., M.M., and M.B., and.; Supervision: M.M., and M.B.

## 5. Material and Methods

### 5.1. Participants and Data

#### 5.1.1. Participants

Participants for MEG analysis were 621 healthy adults (306 females, 315 males) aged 18.5 – 88.9 at the time of neuropsychological assessment. Participants belonged to the Cambridge Center for Ageing and Neuroscience project database ^61^. To evaluate the brain-behavior relationship, we excluded 24 participants with more than three missing cognitive scores among the 8 considered in this study. Additionally, 130 participants without education-related information were excluded from the study (please refer to section 5.1.3). The final sample size comprised 467 participants (237 females and 230 males). Eventual missing scores were imputed by the median value matched to the age decile of the participant.

#### 5.1.2. Magnetoencephalographic (MEG) data

MEG data were collected in a magnetically shielded room in a 306-channel Vectorview MEG system (Elekta Neuromag), 102 magnetometers, and 204 orthogonal planar gradiometers ^93^. A resting-state scan (participants with eyes closed) data ran for 8 min 40 s and sampled at 1kHz (high-pass filter of 0.03 Hz). The MaxFilter 2.2.12 software (Elekta Neuromag, Helsinki, Finland) was used to apply temporal signal space separation (tSSS) ^94^ for noise reduction and offline head motion correction based on the Head-Position Indicator (HPI) coils used to estimate the head position within the MEG helmet.

#### 5.1.3. Cognitive data

Using eight neuropsychological tests, we measured the LP performance, domain-general (DG), and language-specific processes (see Table 1). We also considered education levels as previous work suggested that this factor modulates semantic retrieval ^22^. Education attainment was mapped to a 10-point scale ranging from CSE/GCSE diploma to PhD. To keep homogeneity among participants, we considered the highest degree obtained at age 30.

### 5.2. MEG data processing

#### 5.2.1. Preprocessing and co-registration

MEG analysis steps were performed with the OHBA Software Library (OSL) in Python 3.8.16 following OSL guidelines (https://github.com/OHBA-analysis/osl). Preprocessing began by discarding the first 30 seconds of the raw signals before bandpass filtering using a 5^th^-order IIR Butterworth filter [0.5 - 125 Hz]. Notch filtering was also applied at 50 Hz, 88 Hz, and harmonics (notch width = 2 Hz). This step suppressed interference with power line noise and known artifacts specific to the CamCAN dataset. Subsequently, the data were downsampled to 250 Hz, and automated detection of bad segments and channels was conducted using the generalized-extreme studentized deviate (G-ESD) algorithm ^95^. Further denoising was performed with a FastICA decomposition ^96^, decomposing signals into 64 components and rejecting ECG/EOG-related artifacts. Finally, wrong channels were interpolated from ICA-cleaned data using spherical spline interpolation ^97^.

The resulting MEG data were co-registered to each participant’s structural T1-weighted MR image with an iterative close-point algorithm (ICP) that matched digitized anatomical fiducial points (nasion and bilateral pre-auricular points) to the scalp. The scalp’s surfaces, inner skull, and brain were extracted with FSL’s Brain Extraction Tool (BET).

#### 5.2.2. Source reconstruction and sign flipping

Following OSL guidelines, preprocessed sensor data were bandpass filtered [1 - 45 Hz], and sources were reconstructed onto an 8 mm isotropic dipole grid. This reconstruction was based on a single-shell lead-field model in MNI space and employed a linearly constrained minimum variance (LCMV) scalar beamformer ^98,99^. Continuous time series were extracted and parcellated into 52 regions derived from the HCP-MMP 1.0 atlas ^100^, recently adapted for MEG analysis ^101^. To mitigate source leakage, we applied the symmetric multivariate leakage reduction algorithm ^102^. To address inconsistencies across channels and subjects in source-reconstructed dipole signs, we employed a validated sign-flipping algorithm ^68^.

### 5.3. Hidden Markov Model

This study employed the Time-Delay Embedded variant of the Hidden Markov Model (TDE-HMM) to provide a spatially, spectrally (i.e., defined as a function of frequency) and temporally resolved (i.e., state is active or inactive) description of MEG resting-state neural activity ^68^. Data preparation, model training, and post hoc analysis were carried out with the toolbox *osl-dynamics* ^103^ in Python 3.10.14.

#### 5.3.1. HMM description

The HMM posits that the observed MEG data can be generated from a sequence of hidden brain states whose activations wax and wane over time. The TDE variant specifically models the autocovariance of the signal around time point *t*, that is the covariance between *t* and a time-lagged version of itself *t+L* where *L* defines the time window around *t*. Thus, this embedding may better account for the conduction delay between communicating brain regions ^104^. Mathematically, the embedded space is described using a zero-mean Gaussian distribution ^105^:

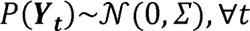

Here, Y_t_ = (y_t-L_,…, y_t_,…, y_t+L_) represents a linear combination of time points *t* for a multichannel time series *y*, and l’ is the multivariate autocovariance matrix.

The TDE-HMM is then defined as:

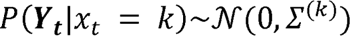

where x_t_ is a hidden variable indicating the state *k* active at time point *t*; and l’^(k)^ is the autocovariance matrix describing the spectral content of the state.

#### 5.3.2. Time-Delay Embedding (TDE)

In line with previous work ^68,106^, we prepared source-space MEG data by embedding *L* = 15 lags evenly distributed across the range −125 Hz to 125 Hz around each time point *t*. Thus, considering 52 parcels, the embedding resulted in 780 channels with 303,810 (780*779/2) parameters to estimate for each state autocovariance matrix. To mitigate overfitting concerns and decrease computational cost, principal component analysis (PCA) was applied to reduce the dimensionality of the embedded space to 104 channels (twice the number of parcels), explaining 65.4% of the variance. The PCA-transformed data were normalized to zero mean and unit variance before model training.

#### 5.3.3. HMM inference

HMM inference was conducted on one GPU NVIDIA A100 Tensor Core 40 Go. We conducted 5 inferences for *K* = 8 states to leverage a low-dimensional data representation that facilitates interpretation and provides a good trade-off between rich and redundant spatio-spectral features across states ^107^. We then carried on with the inference that minimized variational free energy^108^.

### 5.4. Brain states analysis

#### Spectral analysis

As shown in Figure 1, we multiplied the inferred parameters from the model with the unprepared source-space MEG data to retrieve the state time courses. Then, we conducted a multitaper analysis across the 1 – 45 Hz range to extract spectral features^68,109^, resulting in subject- and state-specific spectra.

To enhance interpretability, we determined state-relevant frequency bands in a data-driven manner by running non-negative matrix factorization for each state separately. We asked for a decomposition into 4 frequency bands following the toolbox guidelines, and visually matched each dynamic state’s spectrum to its frequency band of interest (see Figures S3-4, Appendix S2).

#### Spatial Analysis

We mapped brain states onto several resting-state networks. We obtained power maps in the spatial domain for each brain state and region by integrating the corresponding group-averaged spectrum within the corresponding frequency band of interest. The composition of each brain state was then obtained by multiplying the resulting power map with the volumetric overlap between each region and a 12-network atlas^110^ (see Tables S1-2-3, Appendix S3).

### 5.5. Neurocognitive analysis

To examine the relationship between resting-state MEG activity and lexical production (LP) performance across the lifespan, we employed Partial Least Squares (PLS) correlation analysis using the toolbox myPLS (https://github.com/MIPLabCH/myPLS) in MATLAB R2020b. PLS is a statistical technique identifying latent components, capturing coordinated changes between brain state features (*X matrix*) and cognitive (*Y matrix*) performances.

#### 5.5.1. PLS inference

As shown in Figure 1, the covariance matrix (X*Y*^T^*) undergoes singular value decomposition (SVD) to retrieve latent components. Each component is associated with a diagonal set of singular values (S), encoding the amount of shared information, and a set of brain (U) and cognitive (V) saliences, encoding the contribution of each feature. Statistical significance and robustness of saliences were assessed using 10,000 permutations and 1000 bootstrap resamples, respectively. Features with a high bootstrap sampling ratio (BSR ± 3), calculated as the salience weight over its bootstrapped standard deviation, indicate a robust contribution exceeding a 99% confidence interval ^111^.

#### 5.5.2. Data preparation

We examined the temporal and spectral brain state dynamics with three PLS models. As we expected similar latent components, we adjusted all models’ significance threshold at the False Discovery Rate (FDR).

*(i) Model 1: Temporal domain (within states).* For each subject and state, we calculated four metrics. The first two metrics characterize the duration of state activations: Fractional Occupancy (FO), the fraction of total time spent in a state; Lifetime (LT), the mean duration when a state is active. The last two metrics characterize the frequency of state activations: Interval Lifetime (INT), the mean duration between successive activations of the same state; Switching Rate (SR), the mean number of state activations per second.
*(i) Model 2: Temporal domain (between states).* For each subject, we computed the probability that each state gets active after another, resulting in a state-by-state transition probability matrix. To ensure relevancy, we excluded transitions self-transitions in our calculations.
*(i) Model 3: Spectral domain.* We concatenated the power spectra for each subject and state (dimension: subjects *by* states *by* parcels *by* frequencies). We limited our analysis to the top 20 parcels to ensure relevance, showing each state’s most extensive group-level activity.

For the three models, cognitive features were preprocessed in the following steps. First, we regressed out the following covariates: sex, total intracranial volume (TIV), MMSE score, and the number of time points contained in the preprocessed MEG recording of each subject. Second, we quantile-normalized each cognitive score to improve Gaussianity in line with previous work ^112^. Third, we added age-related information as follows:

Given the nonlinear age trajectory reported in MEG studies ^62^, we generated two piecewise polynomials that we included as separate variables in our models. The quadratic term used to generate the polynomials was centered at 55 years old based on converging evidence in different imaging modalities ^19,66,67,113^ and matched to the original scale of the age variable. As shown in Figure 1, the first polynomial function represented the ascending part of an inverted U-shape trajectory and plateaued after age 55. Complementarily, the second polynomial function represented the descending part and accelerated after age 55. If our PLS model maximizes the covariance with a linear age trajectory, then we expect the two functions to have saliences of similar magnitudes but with the opposite sign. In contrast, a quadratic trajectory would yield saliences of similar magnitudes and signs. In sum, this approach allowed us to account, in a data-driven manner, for the optimal age-related trajectory within the subspace defined by linear and quadratic trajectories.

## Data and code availability

Post-hoc data can be downloaded at: https://10.5281/zenodo/12799096. Code is made publicly available at: https://github.com/LPNC-LANG/MEG_CAMCAN_2024

## Supplementary information

**Appendix S1:** Supplementary material Description of the 8 neuropsychological tasks

**Appendix S2:** Supplementary results

- Figure S1-2 – Group-level parcel-averaged spectra
- Figure S3-4 – Spectral decomposition
- Table S1 – Temporal bootstrap sampling ratios for Component 1
- Figure S5 – Spectral bootstrap sampling ratios for Component 2
- Figure S6 – PLS models diagnostics

**Appendix S3:**

- Table S1 – Parcel-by-State map
- Table S2 – Parcel-by-RSN map
- Table S3 – RSN-by-State map

