## Appendix S2 for "Lifespan Oscillatory Dynamics in Lexical Production: A Population-based MEG Resting-State Analysis"

**Appendix S2: Supplementary results**

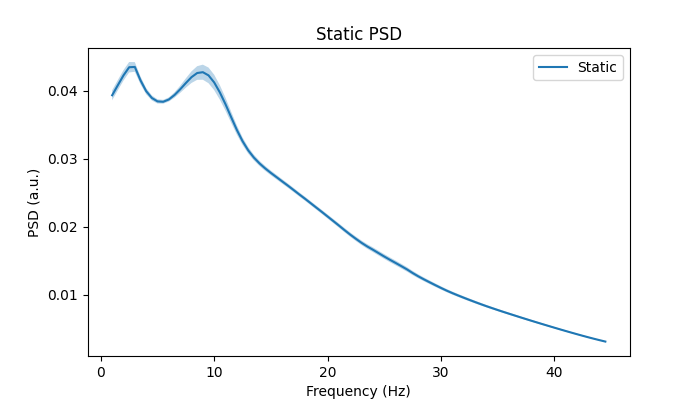

**Figure S1. Group-level parcel-averaged static power spectrum.** This represents the time-averaged spectrum weighted by the brain states’ fractional occupancies. The more time spent in a state, the greater its influence on the static spectrum. *PSD=Power Spectral Density*

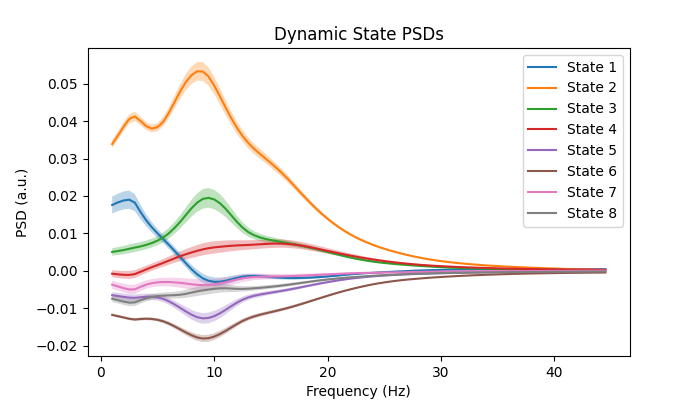

**Figure S2. Group-level parcel-averaged dynamic power spectra.** These spectra were obtained by subtracting the spectrum shown in Figure 1 from each state’s static spectrum, thus reflecting state-specific dynamic activity. *PSD=Power Spectral Density*

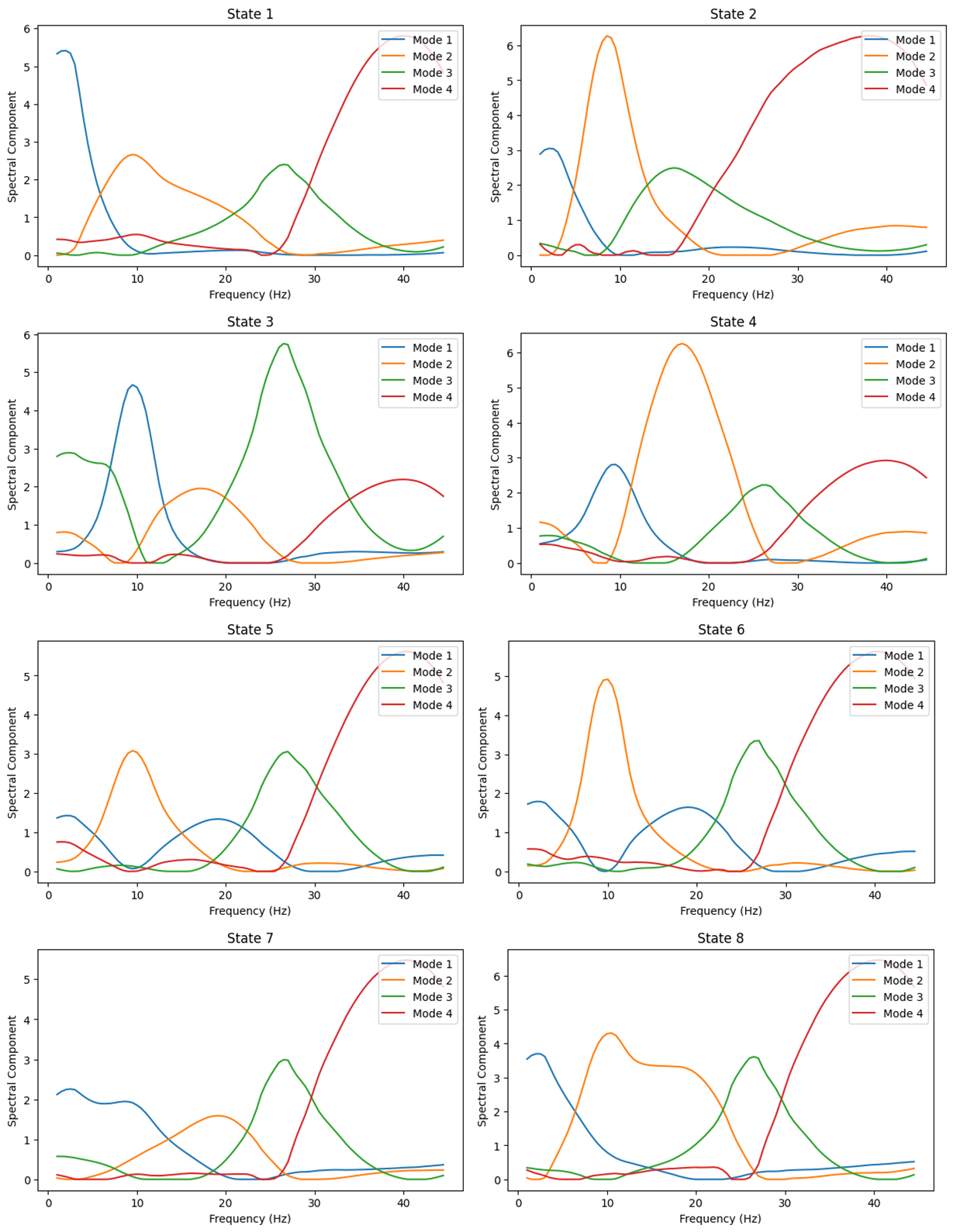

**Figure S3. Spectral decomposition of each state.** This decomposition was obtained by performing Non-Negative Matrix Factorization on the coherence spectra of each state.

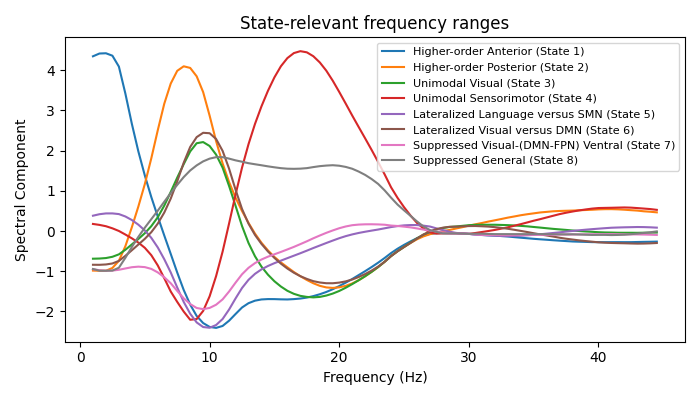
**Figure S4. Group-level state-relevant frequency ranges.** Following the decomposition shown in Figure S3, we matched one of the 4 component visually to the state’s dynamic spectrum shown in Figure S1.

| Brain state  (number – label) | **Duration of state activations** | | **Frequency of state activations** | |
| --- | --- | --- | --- | --- |
|  | ***Fractional Occupancy (FO)*** | ***Mean Lifetime (LT)*** | ***Mean Interval time (INT)*** | ***Switching rate (SR)*** |
| 1- Higher-order Anterior | -12.1 | *ns.* | 10.7 | -17.7 |
| 2- Higher-order Posterior | 11.5 | 7.6 | -9.3 | 7.6 |
| 3- Unimodal Visual | *ns.* | *ns.* | *ns.* | *ns.* |
| 4- Unimodal SMN | *ns.* | 6.9 | 6.6 | -9.6 |
| 5- Lateralized Language vs. SMN | *ns.* | 12.7 | 4.7 | -8.8 |
| 6- Lateralized Visual vs. DMN | 9 | 9.8 | *ns.* | *ns.* |
| 7- Suppressed DMN-FPN-Visual | -16.6 | -9.6 | 10.4 | -12.6 |
| 8- Suppressed General | 11.7 | 8.8 | -9.3 | 6.5 |

**Table S1. Summary of the first latent component's bootstrap sampling ratios (BSR) in the temporal domain.** Only salient (±3) BSRs are reported. *ns. not significant.* This table is illustrated in Figure 4 in the main text.

**Results on the spectral PLS model**

The first latent component explained 58.75% of the total shared variance (p_FDR_ < .001). Spectral changes were linear across the lifespan (*edf* = 1, *F* = 179.2) as opposed to the accelerating cognitive trajectory observed in our study. This suggests that spectral dynamics do not directly reflect cognitive control aspects of lexical production as far as our study is concerned.

However, such low performance could also be explained by methodological concerns: (i) setting an a priori inflection point at age 55 precludes the PLS model from exploring alternative trajectories that may better capture lifespan spectral changes, for example, with an inflection earlier or later in life. (ii) Cross-frequency couplings could better reflect the age-related cognitive trajectory. As shown in Figure S5, we observed oscillatory suppression in the lower and uppermost band (<8 Hz & >30 Hz), which could reflect theta-gamma phase-locking patterns previously associated with attention allocation ^1^, semantic ^2^, sensory and memory processes ^3^ during healthy aging ^4^. More broadly, we found a release in alpha and beta-band power in visuo-occipital areas, which could indicate a trend towards a global inhibition release with aging (see the corresponding channels in yellow in Figure S5 below), together with increased activity in the ~8-25 Hz frequency range over the motor and premotor regions (see the corresponding channels in violet and green) which could indicate enhanced sensorimotor-related information flow across the cortex.

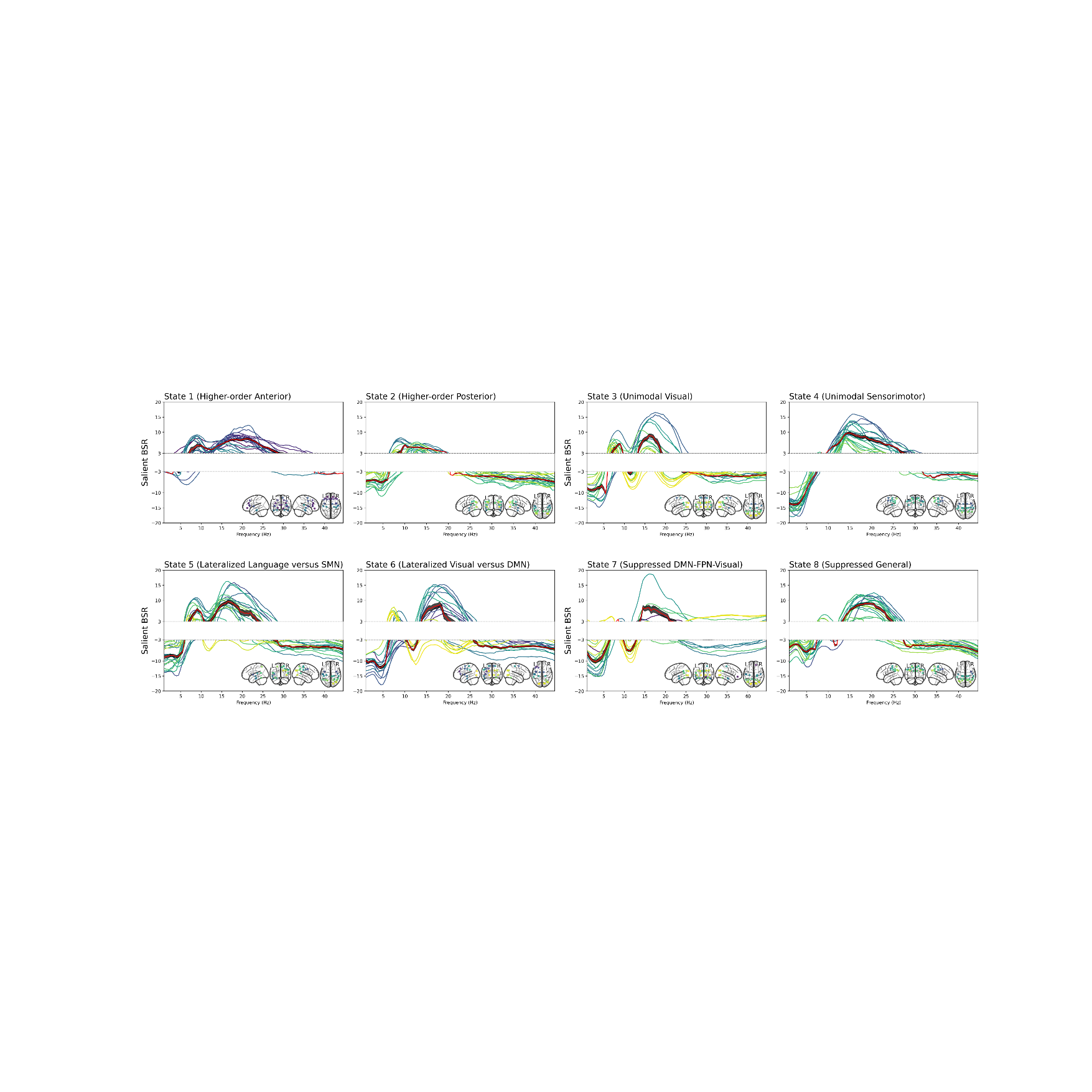
**Figure S5. Bootstrap sampling ratios (BSR) in the spectral domain.** The colors of each channel follow their spatial location (yellow = posterior, green = middle, purple = anterior). Red is the mean salient BSR value; the Grey strip is the standard error. The white band in the range y = -3 to y = 3 masks BSR values below the significance threshold.

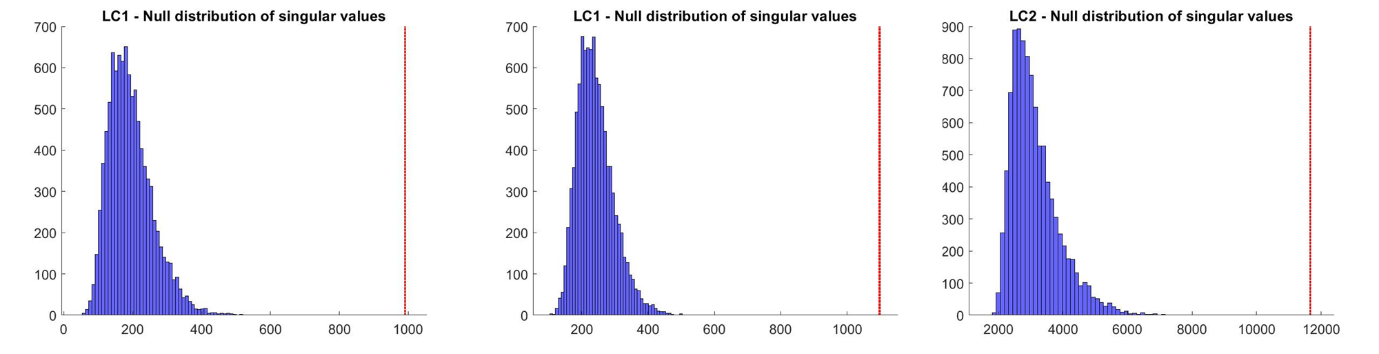

**Figure S6. Partial Least Squares (PLS) model diagnostics.** The null distribution of singular values following 10,000 permutations of the cognitive matrix for the first latent component of the temporal (left) and state-to-state transition (middle) models and the second latent component of the spectral model (left).
